# γ-glutamyl-cysteine is the critical intermediate in glutathione-modulated amelioration of PFOS-neurotoxicity

**DOI:** 10.1101/2025.06.01.657248

**Authors:** Shreesh Raj Sammi, Napissara Boonpraman, Nathan C Kuhn, Jason R. Cannon

## Abstract

Perfluorooctane sulfonate (PFOS) is one of the most prevalent PFAS. Earlier studies using *Caenorhabditis elegans* indicated that glutathione (GSH) ameliorates PFOS-induced neurodegeneration. This study investigates the GSH synthesis pathway to elucidate critical neuroprotective mechanisms. We assessed the effects of GSH precursors, cysteine, glutamic acid, and cysteine + glutamic acid, and of the intermediate γ-glutamyl-cysteine (γ-Glu-Cys), on PFOS neurotoxicity. While cysteine showed slight neuroprotection, other GSH precursors, glutamate, and cysteine + glutamate did not demonstrate beneficial effects. However, the crucial GSH synthesis intermediate, γ-Glu-Cys, conferred neuroprotection comparable to GSH. Notably, no changes were observed in the gene or protein expression of GSH synthesis enzymes in *C. elegans* and SH-SY5Y cells, respectively. Further, testing of the alternative pathway for 5-oxoproline-mediated GSH synthesis revealed that it is not involved in γ-glutamyl-cysteine-mediated neuroprotection. Paradoxically, 5-oxoproline supplementation was neurotoxic, plausibly due to glutamate toxicity. Further, we tested the effect of acute PFOS exposure on mitochondrial complexes I, II, III, and IV. We observed significant inhibition of enzyme activity in mitochondrial complexes II, III, and IV. Next, we tested whether GSH and γ-Glu-Cys could rescue PFOS-induced enzyme inhibition. We found that while both GSH and γ-Glu-Cys could rescue enzyme inhibition in complex III, the rescuing effect was altogether absent for complex IV. Overall, our findings confirmed that both GSH and γ-Glu-Cys can rescue PFOS-induced neurodegeneration and mitochondrial enzyme affliction. This study effectively uncovers a novel mechanism that addresses existing knowledge gaps pertaining to the role of the GSH pathway in curtailing PFOS neurotoxicity.

**Graphical Abstract:** Graphical Abstract γ-glutamyl-cysteine is the critical intermediate in glutathione-modulated amelioration of PFOS-neurotoxicity
Perfluorooctanesulfonate (PFOS) leads to dopaminergic cell loss in *C. elegans*. Glutathione (GSH) confers neuroprotection against PFOS neurotoxicity. Further investigation of the critical components involved in GSH biosynthesis identified that while precursors of GSH, Cysteine and glutamate fail to confer neuroprotection, only γ-glutamyl-cysteine alleviates neuronal loss similar to GSH.

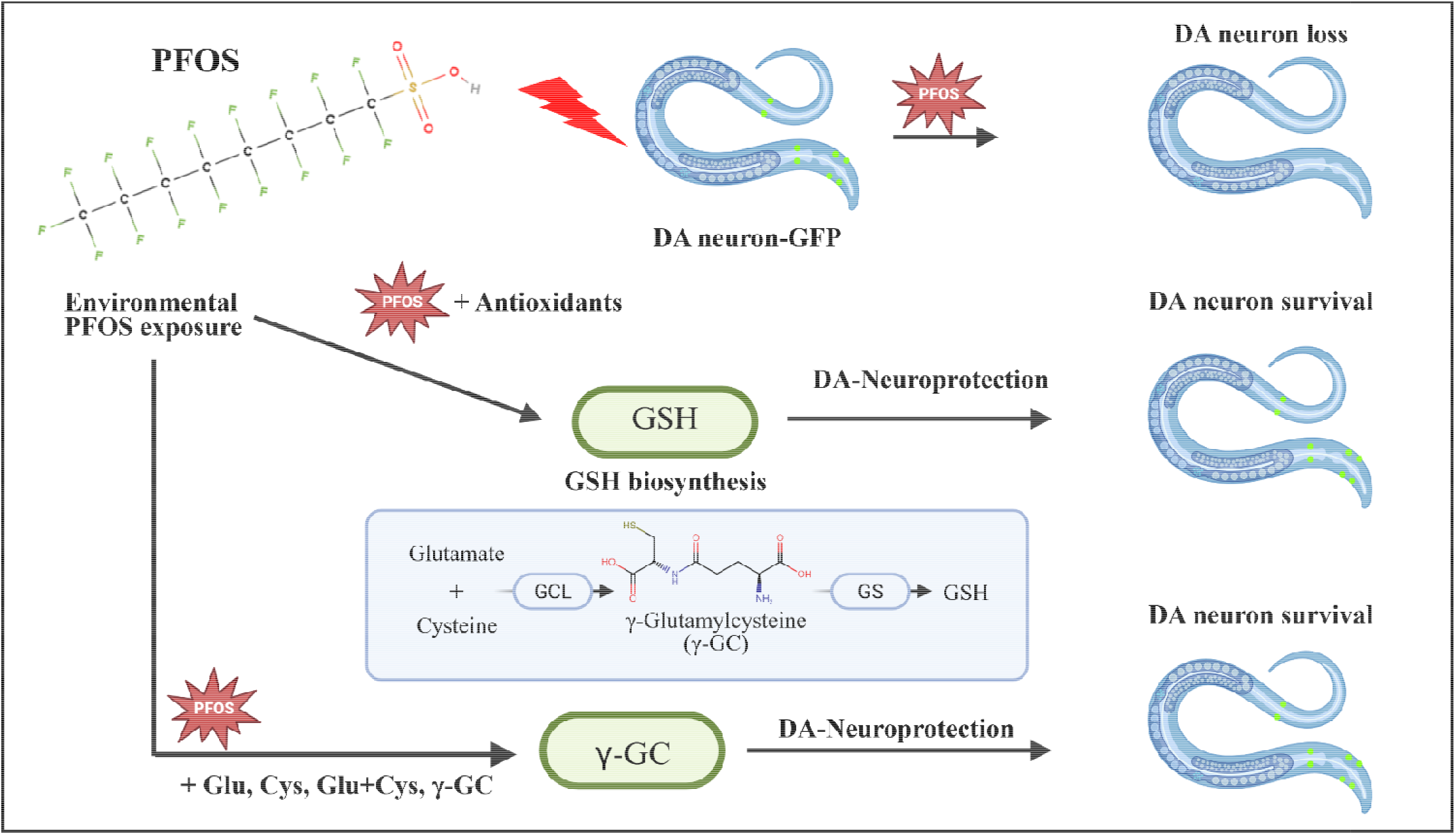

## Introduction

Per- and polyfluoroalkyl substances (PFAS) are anthropogenic chemicals that are used in several consumer and industrial applications such as manufacture of textiles, carpets, upholstery, leather, waxes, food packaging, adhesives, paints, polishes, cosmetics, aviation hydraulic fluids, firefighting foams, and papers [1]. Environmental persistence is due to their resistance to degradation[2], owing to the strong carbon-fluorine bond[3] posing a significant risk to the environment, wildlife and human health. PFAS can be classified as legacy and emerging with respect to their structure, legacy being long-chain and emerging as short chain and branched. Despite phase-out, perfluorooctanesulfonate (PFOS) and perfluorooctanoic acid (PFOA) are one of the most prominent legacy PFASs[4]. Amongst the two PFOS has the longest half-life in human body (∼3.4 to 5.7 years in serum)[5].

Recent data from a French cohort investigating the PFAS concentration in cerebrospinal fluid samples from patients living away from fluorochemical industry has shown that while the emergent PFAS tend to be eliminated, PFOS persists, and the concentration increases with age[6]. Hence, it is important to investigate PFOS with respect to its neurotoxicological aspects. Previous studies from our group using *C. elegans* models have indicated that PFOS is neurotoxic, particularly to the dopaminergic neurons. Importantly, while numerous glutathione relevant antioxidants, including N-acetyl cysteine failed to ameliorate neurotoxicity, GSH alone was found to be neuroprotective. Further, mitochondria were found to be primary neurotoxic targets, exhibiting dysfunction and damage at far lower doses than those that induced cell death or other markers of neurotoxicity [7]. Given that GSH is the principal mitochondrial antioxidant[8-10]It is critical to investigate the role of the GSH pathway and its implications on GSH-led mitigation of PFOS neurotoxicity. Thus, this study aimed to identify the critical intermediate in the GSH pathway and specific neuroprotective mechanisms to PFOS exposure.

### Experimental Procedures

#### Culture and maintenance of Strains

*Caenorhabditis elegans* strains, N2 and BZ555 (egls1[dat-1p::GFP]), and *Escherichia coli* OP50 were procured from the Caenorhabditis Genetics Center (University of Minnesota, Minnesota), grown on Nematode growth medium (NGM), and cultured at 22°C. A synchronized population of worms was obtained by sodium hypochlorite treatment. Embryos were incubated overnight at 15°C in M9 buffer to obtain L1 worms.

#### Treatment of worms

L1-stage worms were treated with different concentrations of PFOS as described in K medium as described previously[11,12]. 10,000-ppm stock of PFOS was prepared in methanol and diluted further to doses ranging from 25 to 75 ppm. Approximately 175 L1-stage worms were treated for 72 hrs. in 12-well plates. The concentration of methanol was adjusted accordingly for each experimental group, with the final concentration being 0.75%. On the basis of results obtained from this study and previous research[7], co-exposures were conducted at 75 ppm, PFOS.

To investigate the effect of precursors and intermediates of GSH pathways, cysteine (Cys), Glutamate (Glu), and γ-glutamyl-cysteine (γ-Glu-Cys) worms were co-exposed with PFOS 75 ppm within the dose range 0.125 to 2 mM for 72 hrs. The dose paradigm is based on GSH doses used earlier[7]. 100 mM stock of 5-Oxoproline (5-OP) was dissolved in K media complete. BZ555 worms were treated with 5 and 10 mM 5-OP. Co-exposure studies with PFOS 75 ppm were conducted at dose range: 0.125, 0.25, and 0.5 mM 5-OP.

#### Neurodegeneration Assay

Neuropathological effects of PFOS on dopaminergic neurons were studied by treating L1 stage worms for 72 hrs at 22 °C. Neurodegeneration was scored as described previously[7,12,13]. Briefly, treated worms were washed three times using M9 buffer and anesthetized using 10 µL of 100 mM sodium azide. Counting of neurons was done for all neuron types i.e., Cephalic sensilla (CEP), Anterior deirid (ADE), and posterior deirid (PDE), using a FITC filter. *C. elegans* have eight DA neurons, 4 CEP, 2 ADE, and 2 PDE [14]. The percentage of intact neurons (PIN) was calculated for a minimum of 20 worms per group (independent replicates). IC50 values for GSH and γ-Glu-Cys were calculated by normalizing each set of experiments with respect to the control and multiplying by 100 to generate percentages. Finally, toxicity inhibition was calculated by subtracting the obtained values with 100.

#### RNA Isolation, cDNA Synthesis, and Quantitative Real-Time PCR

Total RNA was extracted from worms exposed to PFOS 50 ppm and 75 ppm for 72 hrs using RNAzol reagent (Molecular Research Centre Inc.) as per manufacturer’s instructions. In brief, worms were washed with nuclease free water and homogenized in 200 µL of RNAzol, followed by incubation at room temperature for 10 minutes. The homogenate was centrifuged at 13000 rpm for 10 minutes at 4°C. Supernatant was added to 100 µL of nuclease-free water. Centrifuge tubes were vortexed for 15s and were allowed to sit for 15 minutes at room temperature. RNA was precipitated using equal volume of isopropanol and allowed to sit for 15 minutes at room temperature. Finally, the RNA was pelleted by centrifugation at 13000 rpm for 10 minutes at 4°C. RNA pellet was washed twice using 75% ethanol by centrifugation at 6500 rpm for 5 minutes at 4°C. RNA samples were quantified and assessed for quality using nanodrop. cDNA synthesis was done using 1 µg of RNA in a thermal cycler using RevertAid H minus First Strand cDNA synthesis Kit (Thermo Scientific) manufacturer’s protocol. qRT-PCR studies were done using BioRad CFX 96. Differential expression was calculated by 2^-ΔΔCT^ method[15]. *Gpd-1* was used as internal control and used to calculate fold change in mRNA expression. Primers were procured from Integrated DNA technologies, with details as shown in table 1 (orientation of oligonucleotides: 5’ to 3’).

**Table 1:**
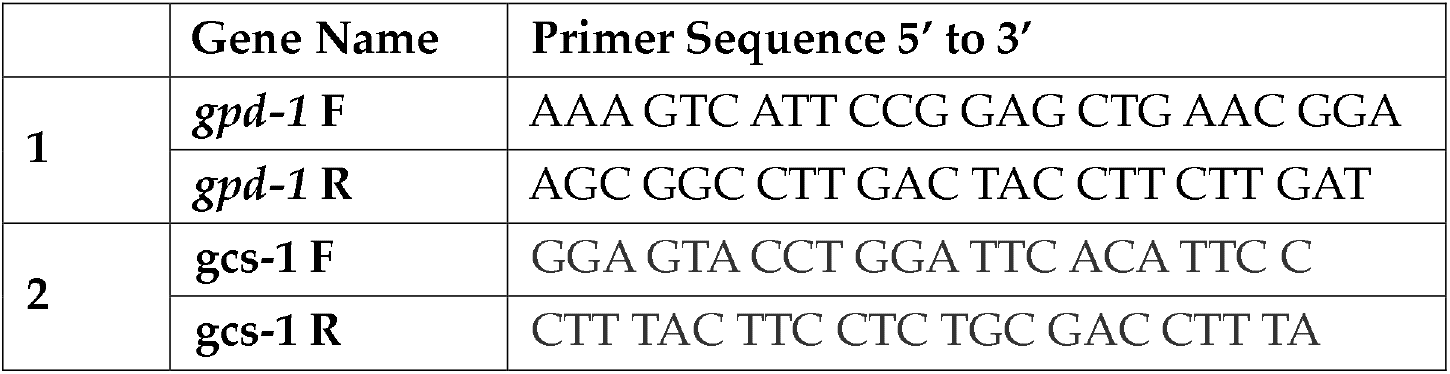
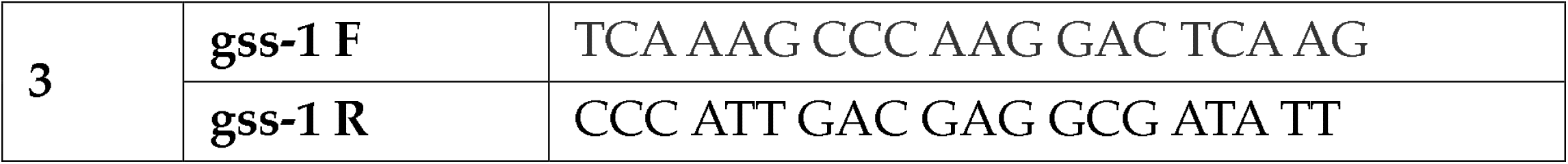
Sequence of the primers used.

#### SH-SY5Y cells culture and treatment

SH-SY5Y cells were procured from ATCC (American Type Culture Collection, Manassas, VA). Cells were cultured using Basic Growth Media (BGM) as described previously[16]. 90% Confluent cells were treated with different concentrations of PFOS (25, 50, and 75 ppm) for a period of 24 hrs.

#### Immunoblotting

Total protein was harvested from SH-SY5Y cells through sonication and quantified using Pierce BCA protein assay kit per manufacturer’s protocol. 80 µg of protein was loaded per lane in 4-15% precast protein gels (Bio-Rad Laboratories, Inc.) and run using a Bio-Rad Mini Protean electrophoresis unit. Protein was transferred to nitrocellulose membrane using Bio-Rad semidry transfer unit at 10 V for 30 minutes. Membranes were blocked using 5% BSA in PBS for 1 hr. and probed with antibodies - β-actin, Glutathione cysteine ligase (GCL), and Glutathione synthetase (GS) (with blocking solution; overnight incubation on rocking at 4°C). All antibodies were procured from Abcam Inc., Cambridge, U.K. Secondary antibodies were procured from Li Cor Biosciences, Lincoln, NE. PBST 0.05% was used for washing (∼5 minutes on orbital shaking) the membrane, and antibody incubation was done in PBST 0.01%. Membranes were visualized using Li-Cor Odyssey clx, and protein bands were quantified using Image Studio, Version 5.5.

#### Total GSH estimation

Total GSH was estimated for SH-SY5Y cells, treated with PFOS (25, 50, and 75 ppm) for 24 hrs. using Kerafast RealThiol AM Ester Glutathione detection kit as per the manufacturer’s protocol and Jiang et al., 2020[17]. Briefly, post-treatment with PFOS, cell culture media was replaced with the imaging buffer and stained at room temperature for 15 minutes. Fluorescence was read using Synergy H1 microplate reader (Biotek Instruments, Inc., Winooski, VT) using green and blue channels, 488 nm and 405 nm excitation, respectively.

### Animals

Four- to six-month-old male Fischer 344 rats (CDF Strain Code 002; total n = 11) were purchased from Charles River Laboratories. Rat liver was used as a source of mitochondria because it is an ideal organ for bulk extraction, enabling rapid isolation of large quantities of intact mitochondria[18]. Mitochondria were isolated from two independent cohorts of animals. The initial cohort consisted of three rats housed at Purdue University, and the experiment subsequently continued using a second cohort of eight rats housed at Michigan State University (MSU). Rats were maintained under standard laboratory conditions with a 12 h light/dark cycle and provided ad libitum access to food and water. All animal procedures were conducted in accordance with the protocols and were approved by the Institutional Animal Care and Use Committees (IACUC) at the respective institutions. Rats were euthanized by decapitation, and livers were rapidly excised. Mitochondria were immediately isolated as described[19] and stored at −80°C until further use.

### Mitochondrial Enzyme Activity Assay

Mitochondrial respiratory chain enzyme activities (Complexes I–IV) were measured following the spectrophotometric method previously described in our study[12], which was adapted from Spinazzi et al., 2012[19] . Isolated mitochondria were acutely co-exposed to PFOS (50 and 75 ppm), with 0.25 mM GSH or 0.25 mM γ-glutamyl-cysteine (γ-Glu-Cys), for 15 minutes. Following substrate addition, absorbance was recorded every 10 s for 3 min. Enzyme activity (EA) was calculated using the following equation: EA (nmol min^-1^ mg^-1^) = (Δ Absorbance/min × 1,000)/[(extinction coefficient × sample volume (mL)) × sample protein concentration (mg mL^-1^)]. Relative enzyme activity was expressed after normalization to the vehicle control.

#### Statistical Analysis

Statistical analyses were performed using GraphPad Prism version 10 (GraphPad Software, La Jolla, CA, USA). All experiments were performed with at least three independent biological replicates. Data were normalized to vehicle control unless otherwise specified. Statistical significance was determined using one-way or two-way analysis of variance (ANOVA), followed by Dunnett’s multiple comparisons test, as appropriate. Data are presented as the mean ± SD, and differences were considered statistically significant at *p* < 0.05.

## Results

### PFOS produces dose-dependent neurodegeneration

In confirmation of our previous studies showing concentration dependent neuronal loss in response to PFOS[7], we re-assessed the effect of PFOS on dopaminergic (DA) cell loss. A significant decrease in percentage of intact neurons (PIN) was observed in worms treated with 50 ppm (76.87 ± 1.083, *p* < 0.0001) and 75 ppm (54.167 ± 3.554, *p* < 0.0001) in comparison to the control (Figure 1A, 1B). EC50 value of PFOS was found to be ∼157.3 µM. These results were in concurrence with our previous findings.

**Fig. 1:**
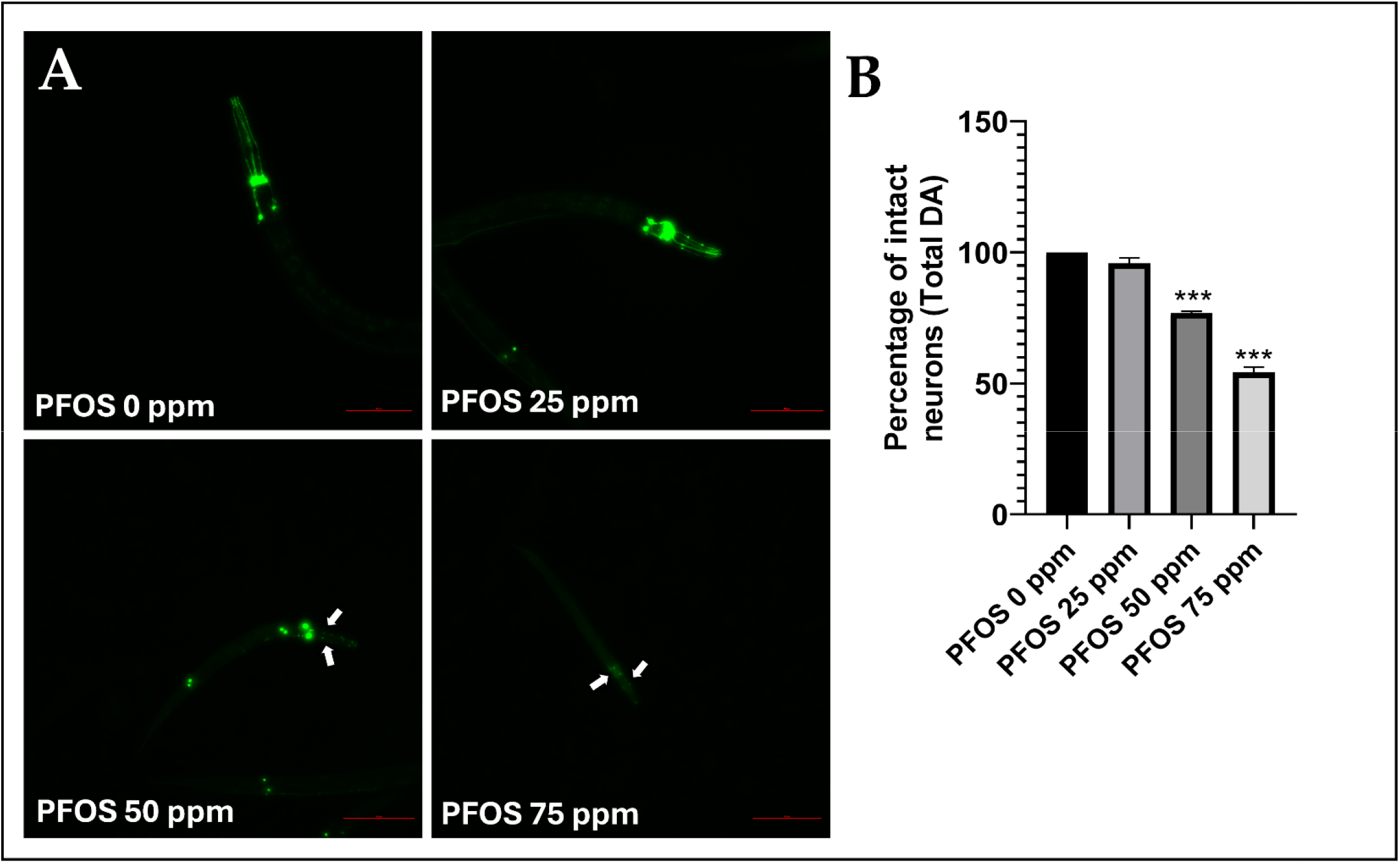
PFOS treatment results in dopaminergic cell loss. Treatment with PFOS led to a decrease in the percentage of intact neurons (PIN) in a concentration-dependent manner. Representative fluorescence micrographs (A) and quantification of the percentage of intact neurons (PIN) (B) following PFOS exposure. L1 worms were exposed to PFOS for 72 h, resulting in a concentration-dependent decrease in PIN. Data are presented as the mean ± SD (n = 3). Statistical analysis was performed using one-way ANOVA followed by Dunnett’s multiple comparisons test. ***p < 0.001. White arrowheads indicate dopaminergic neuron loss. Scale bar = 100 μm.

### Precursors of GSH do not alter PFOS neurotoxicity

Our previous research showed that amongst the several antioxidants tested, only GSH could curtail the PFOS neurotoxicity[7]. Thus, we aimed to discover the role of GSH biosynthesis pathway, testing the precursors of the GSH (Figure 2A) as ameliorators of PFOS neurotoxicity. BZ555 worms were co-exposed with PFOS and Cys, Glu, and Cys + Glu (dose range: 0 to 2 mM). All the co-exposures (Figure 2B, 2C, 2D, 2E) except Cys 0.125 mM (69.16 ± 4.161, p = 0.0157), in comparison to the PFOS 75 ppm (54.583 ± 1.301) as shown in Figure 2B, 2C.

**Fig. 2:**
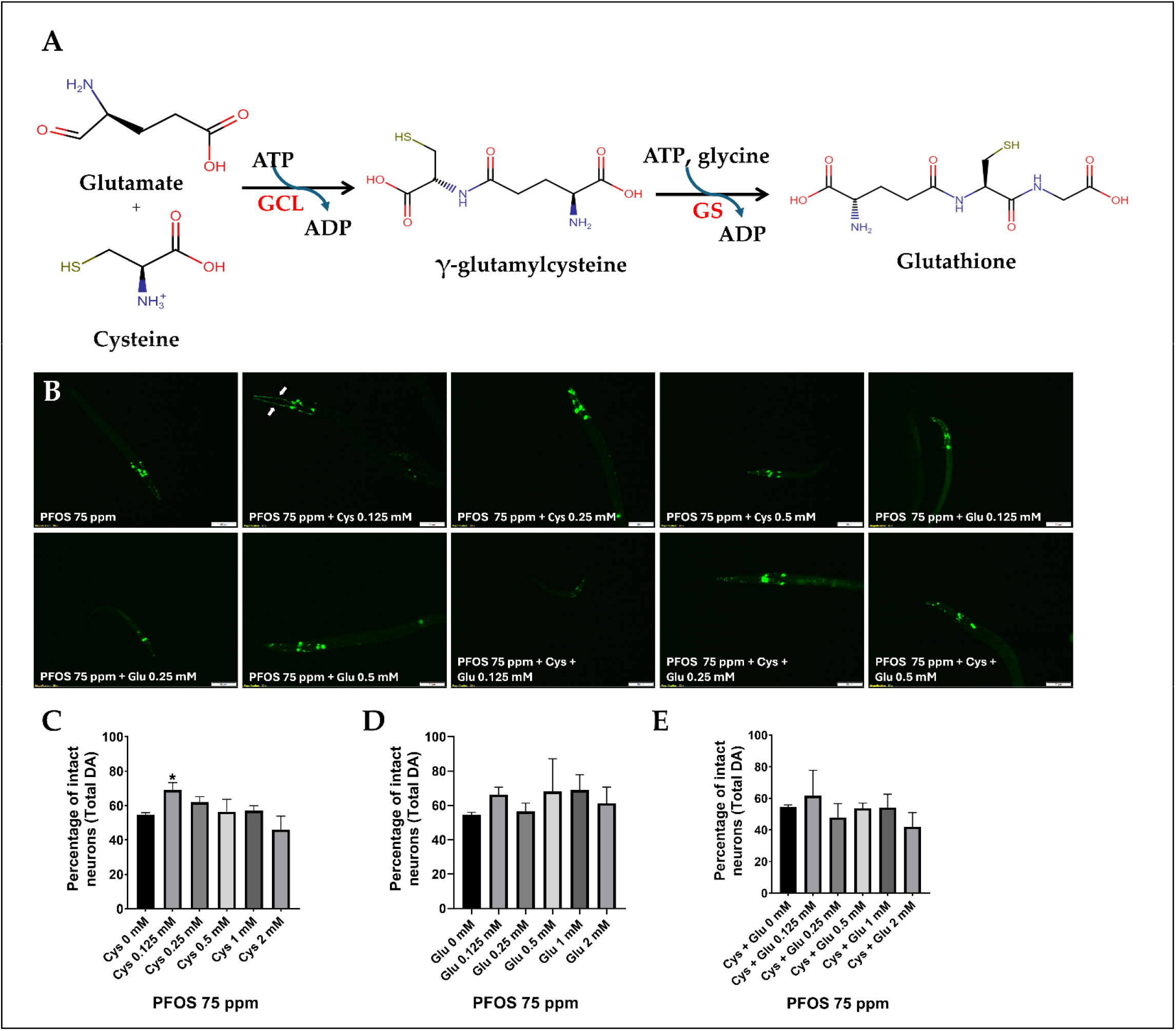
Effect of GSH biosynthesis precursors on PFOS led neurodegeneration. Mechanism of GSH synthesis (A). Representative image showing effect of γ-glutamyl-cysteine on PFOS-led neurodegeneration (B). Co-treatment with cysteine led to slight amelioration at 0.125 mM of PFOS-mediated neurodegeneration (C). Co-treatment with Glutamic acid was devoid of any effect on PFOS-led neurodegeneration (D). Co-treatment with cysteine and glutamic acid combined did not show any significant effect (E). L1 worms were treated with PFOS and GSH precursors for 72 h. Data presented as mean ± SD (n=3). Data was analyzed using one-way ANOVA followed by Dunnett’s post hoc test. \**p* <0.05, \*\**p* <0.005, and \*\*\**p* <0.001. White arrowheads indicate dopaminergic neuron recovery. Scale bar represents 50 μm.

### γ-glutamyl-cysteine ameliorates PFOS neurotoxicity

Given the inability of GSH precursors to mitigate PFOS neurotoxicity, we investigated the key intermediate, γ-glutamyl-cysteine. We observed a significant amelioration of neurodegeneration in worms co-exposed with γ-Glu-Cys 0.125 mM (68.906 ± 2.414, *p* = 0.0151), 0.25 mM (68.281 ± 5.737, *p* = 0.0191), and 0.5 mM (69.688 ± 4.034, *p* = 0.0112) in comparison to PFOS 75 ppm (50.156 ± 4.203) alone (Figure 3A, and 3B). Notably, this effect was similar to GSH conferred neuroprotection (IC50 for GSH = 0.1149 and IC50 for γ-Glu-Cys = 0.07563) as shown previously[7]. The percentage inhibition of neurotoxicity between GSH and γ-Glu-Cys is shown in Figure 3D. These results confirmed that γ-Glu-Cys is the critical intermediate in the GSH pathway, contributing to the mitigation of PFOS neurotoxicity.

**Fig. 3:**
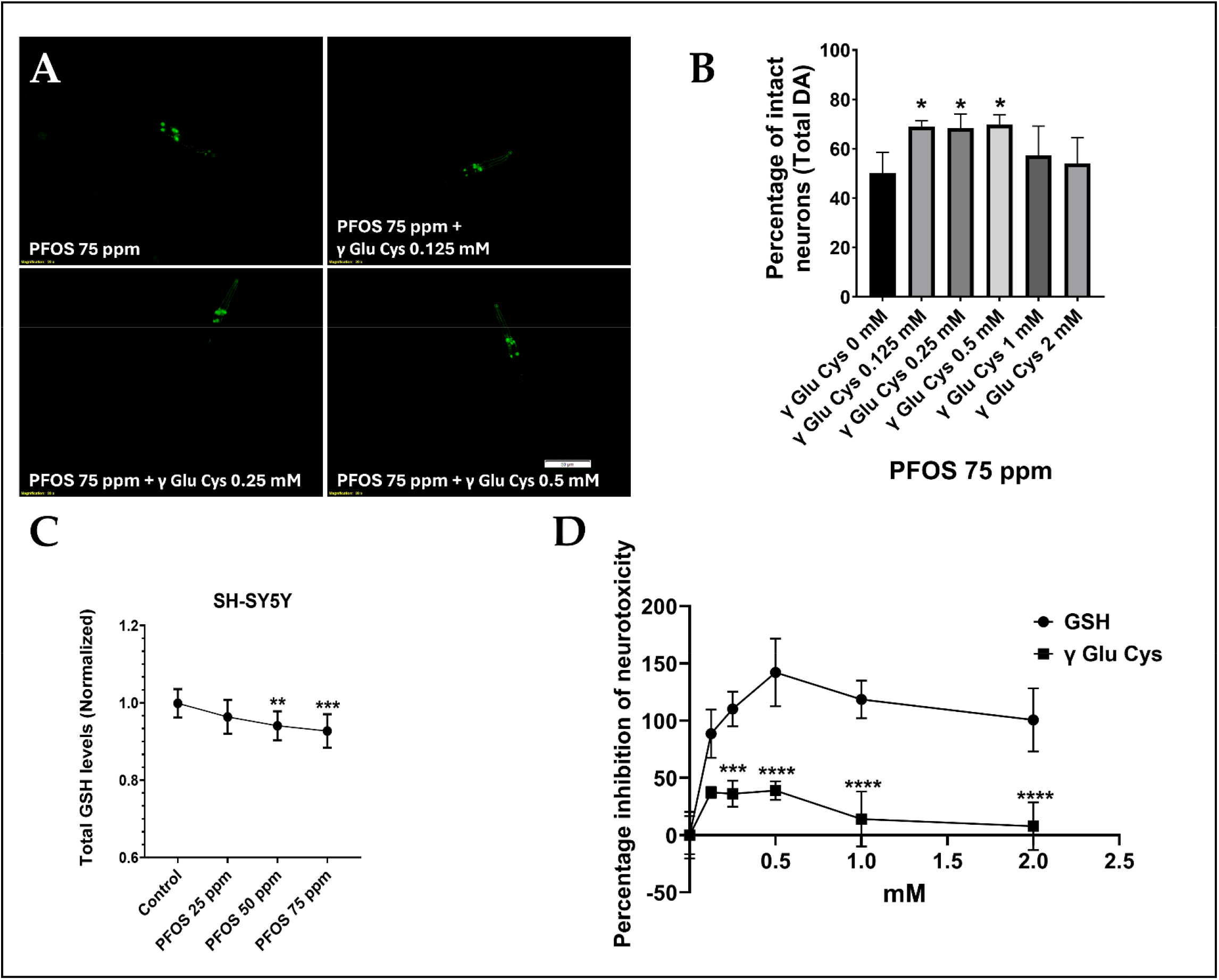
γ-Glutamyl-cysteine alleviates PFOS neurotoxicity. Co-treatment with γ-glutamyl-cysteine significantly attenuated PFOS-induced dopaminergic neurotoxicity at concentrations ranging from 0.125 to 0.5 mM. These findings identified γ-glutamyl-cysteine as a critical intermediate in the GSH biosynthesis pathway, with respect to PFOS-led neurodegeneration (A, B). A decrease in total GSH levels upon treatment wi h PFOS was also observed in SH-SY5Y cells (C). Worms were treated for 72 h.; SH-SY5Y cells were treated with PFOS for 24 h. at 90% confluency. Comparison of the percentage inhibition of neurotoxicity GSH vs. γ-Glu-Cys showed that GSH was relatively more neuroprotective (D). Data presented as mean ± SD (n =3). Statistical analysis was performed using one-way ANOVA followed by Dunnett’s multiple comparisons test or two-way ANOVA, as appropriate. \**p* <0.05, \*\*\**p* <0.001, and \*\*\*\**p* <0.0001. Scale bar = 50 μm.

### PFOS modestly alters glutamyl cysteine ligase gene expression

To further investigate the effect of PFOS on GSH synthesis pathway, we tested the effect of PFOS on the enzymes related to GSH biosynthesis. qPCR was performed for the genes *gcs-1*/GCL (Glutamate cysteine ligase), and *gss-1*/GS (glutathione synthetase). No effect on mRNA expression of *gss-1* (Figure 4A) was observed, while a slight downregulation of *gcs-1* was observed in worms exposed to PFOS 50 ppm (Fold change: 0.616 ± 0.136, *p* = 0.0221) (Figure 4A).

**Fig. 4:**
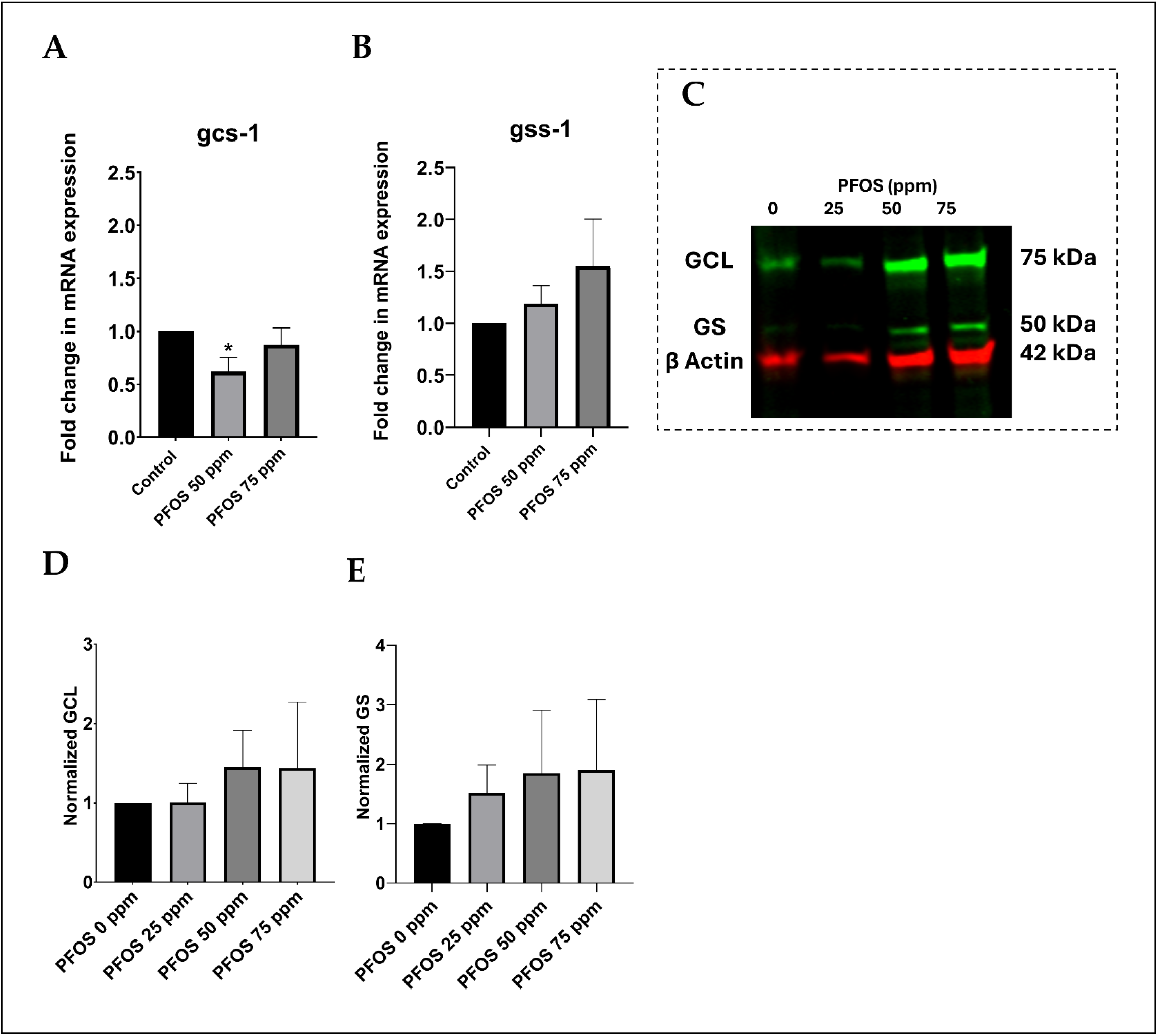
PFOS neurotoxicity does not alter expression of enzymes involved in GSH synthesis. PFOS slightly downregulated *gcs-1/GCL* mRNA expression (A), whereas *gss-1/GS* mRNA expression remained unaltered (B). Representative immunoblot showing GCL and GS protein expression in SH-SY5Y cells (C). Quantification of GCL and GS protein levels showed a slight but nonsignificant increase following PFOS treatment (D, E). Data are presented as the mean ± SD. Statistical analysis was performed using one-way ANOVA followed by Dunnett’s multiple comparisons test. \**p* <0.05 (n = 3 for A and B; n = 4 for D and E).

### PFOS does not alter expression of enzymes involved in GSH synthesis

Upon testing the effect of PFOS on the expression of enzymes involved in GSH synthesis, we further investigated the enzymes using immunoblotting. While there was a slight increase in expression of enzymes, GCL and GS in SH-SY5Y cells were observed, it was statistically insignificant (Figure 4C, 4D, and 4E).

### 5-Oxoproline exacerbated PFOS-led neuronal loss

We further addressed the fate of γ-Glu-Cys by testing the effect on alternate pathways. Excess γ-Glu-Cys is acted upon by the enzyme, γ-GlutamylCycloTransferase producing cysteine and 5-Oxoproline[20]. 5-Oxoproline (5-OP) is catalyzed by the enzyme 5-oxoprolinase to yield glutamate[21]. Glutamate is also known to be neurotoxic[22,23]. So first, we tested the neurotoxicity of higher doses of 5-OP and used 1/40 to 1/10 of the dose to be co-administered with PFOS at 75 ppm.

We observed a complete loss of dopaminergic neurons, both cell bodies and dendrites, at 5 mM OP (0.625 ± 1.0836, *p* = 0.0001), and 10 mM OP (0.000 ± 0.000, *p* = 0.0001) in comparison to 100% intact neurons in control worms (Fig 5A, B). Next, we tested the effect of 0.125, 0.25, and 0.5 mM 5-OP on worms exposed to PFOS 75 ppm. As expected, a significant decrease in PIN was observed in case of worms co-exposed to 0.25 mM 5-OP (31.250 ± 9.615, *p* = 0.0035), and 0.5mM 5-OP (21.406 ± 4.878, *p* = 0.0003) in comparison to PFOS 75 ppm (58.906 ± 6.381) (Fig 5C, D). These results confirm that the observed neuroprotective effect of γ-Glu-Cys was due to conversion of γ-Glu-Cys to GSH and does not involve alternative pathway leading to 5-OP synthesis.

**Fig. 5:**
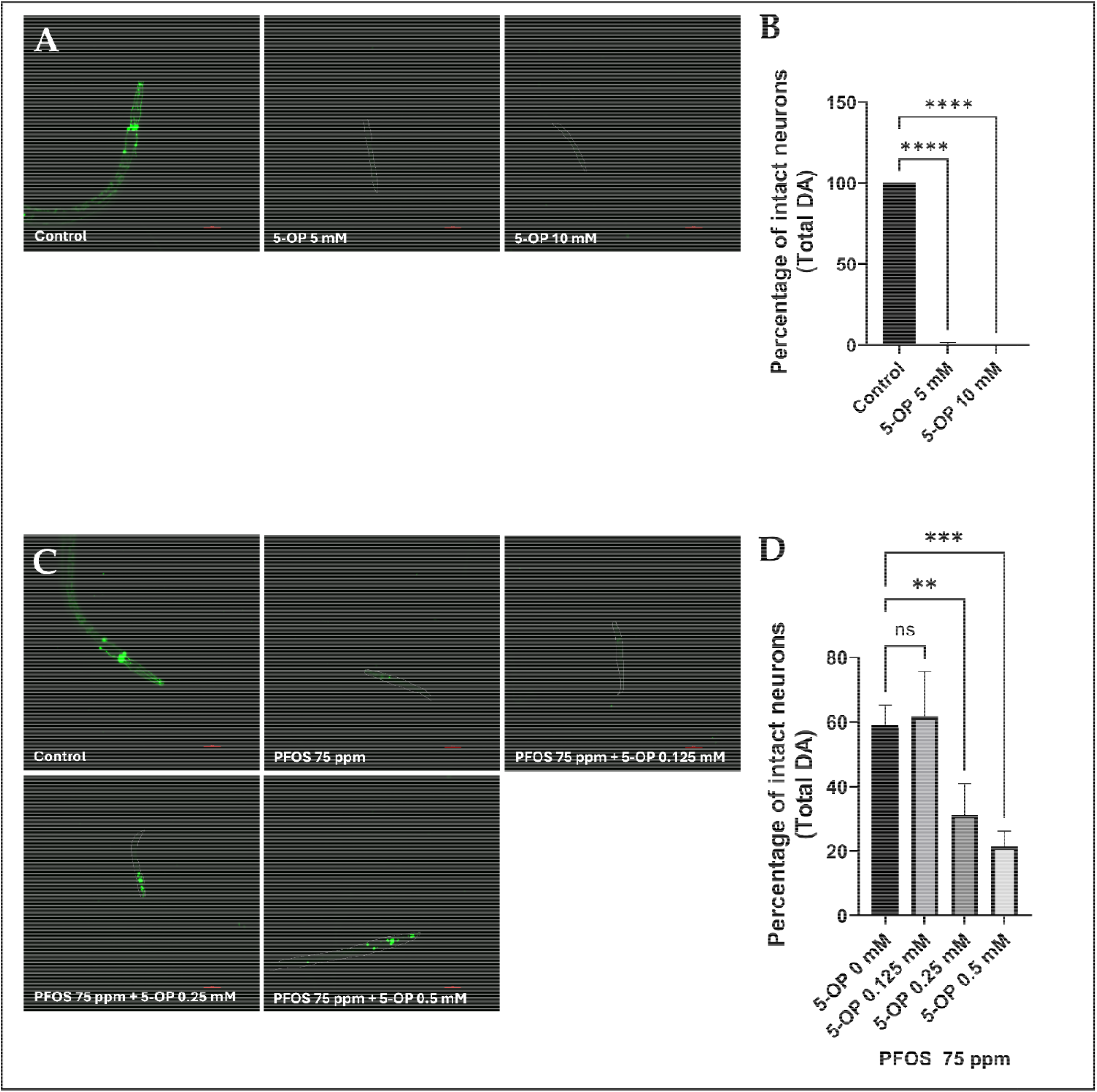
5-Oxoproline treatment exacerbates PFOS-led neurodegeneration. 5-oxoproline (5-OP), a downstream intermediate in the glutathione (GSH) cycle and a precursor of glutamate. Representative fluorescence micrographs (A) and quantification of the percentage of intact dopaminergic neurons (B) following treatment with 5-OP alone. Exposure to 5 or 10 mM 5-OP resulted in complete dopaminergic neuron loss. Representative fluorescence micrographs (C) and quantification (D) showing the effects of co-treatment with PFOS (75 ppm) and 5-OP (0.125–0.5 mM). Co-treatment with 0.5 mM 5-OP significantly exacerbated PFOS-induced dopaminergic neurodegeneration, resulting in a decreased percentage of intact neurons. L1 worms were exposed to 5-OP alone or co-treated with PFOS and 5-OP for 72 h. Data are presented as the mean ± SD. Statistical analysis was performed using one-way ANOVA followed by Dunnett’s multiple comparisons test. \**p*< 0.05, \*\**p* < 0.01, and \*\*\**p* < 0.001 (n = 3 for A and B; n = 4 for C and D). Scale bar = 100 μm.

### PFOS inhibits mitochondrial complex II, III, and IV

Our earlier studies identified mitochondria as one of the primary targets of PFOS[7]. Hence, we tested the effect of PFOS on mitochondrial enzyme activity for complexes I to IV, by acutely exposing isolated mitochondria to different doses of PFOS (0 to 250 ppm) for 15 minutes. While no effect on Complex I enzyme activity was observed, a significant inhibition in normalized enzyme activity of complex II was observed at doses, 125 ppm (0.781 ± 0.056,*p* = 0.0148) and 250 ppm (0.754 ± 0.032, *p* = 0.0065) in comparison to control (1.000 ± 0.000) (Figure 6B). A higher effect was observed in case of complex III, exhibiting inhibition at doses, 50 ppm (0.480 ± 0.216, *p* = 0.0050), 125 ppm (0.0.16 ± 0.190, *p* < 0.0001), and 250 ppm (0.182 ± 0.056, *p* < 0.0001) in comparison to respective control (1.000 ± 0.000) (Figure 6C).

**Fig. 6:**
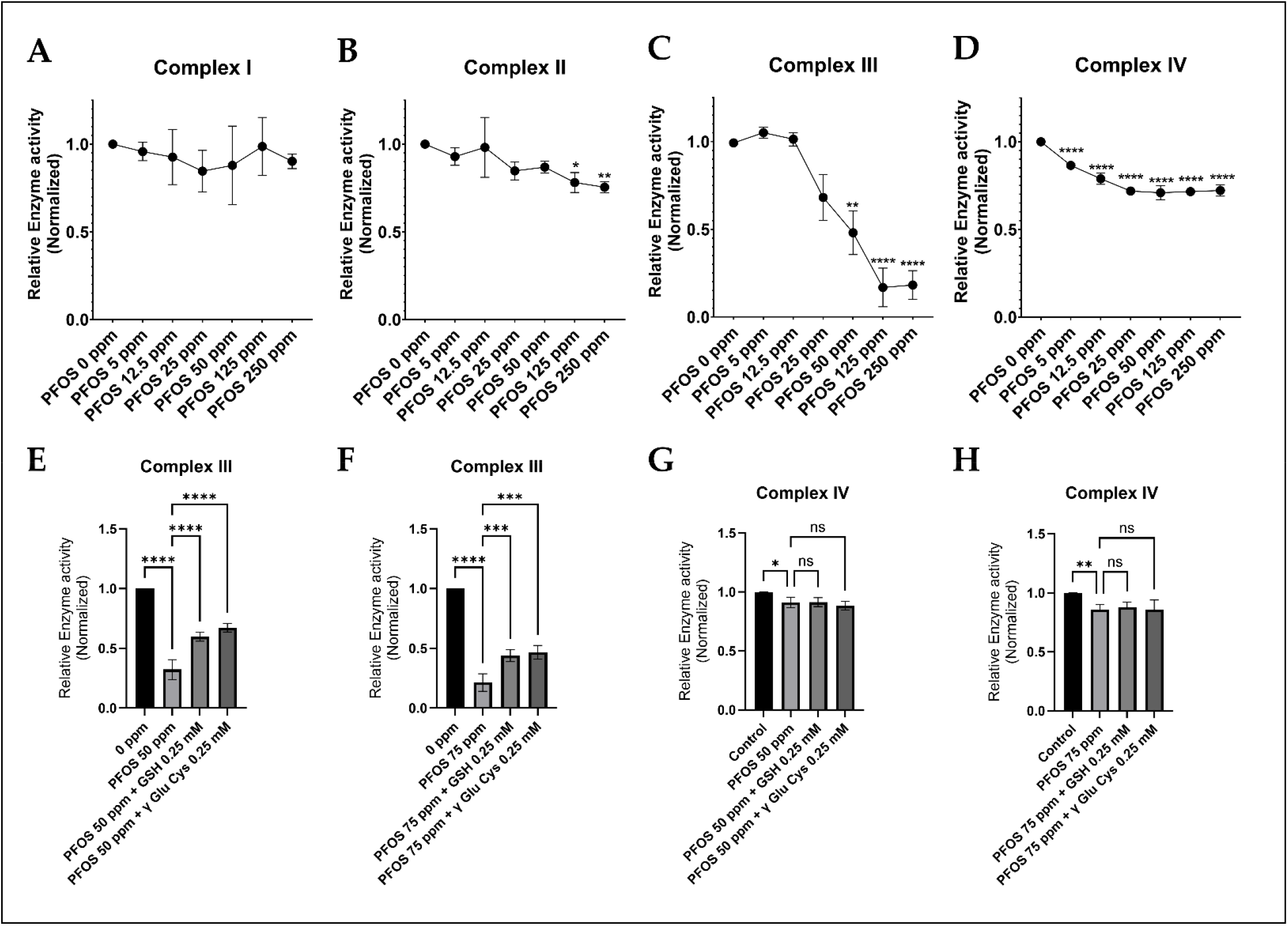
Acute PFOS exposure inhibits mitochondrial enzyme activity. Isolated mitochondria were acutely exposed to PFOS for 15 minutes and assessed for the enzyme activity for the complex I, II, III, and IV. While complex I was devoid of any effect (A), a significant dose dependent inhibition was observed in the enzyme activity of complex II (B), Complex III (C), and complex IV (D). Further, the ameliorative effect of GSH and γ-Glu-Cys was tested on the PFOS led inhibition of Complex III, and IV. While both GSH and γ-Glu-Cys exhibited alleviation of complex III inhibition at PFOS 50 ppm (E) and PFOS 75 ppm (F), the rescuing effect was absent in case of PFOS led complex IV inhibition (G, H). Data are presented as the mean ± SD. Statistical analysis was performed using one-way ANOVA followed by Dunnett’s multiple comparisons test. \**p*< 0.05, \*\**p* < 0.01, and \*\*\**p* < 0.001 (n = 3 for A, B, C, and D; n = 4 for E, F, G and H).

**Fig. 7:**
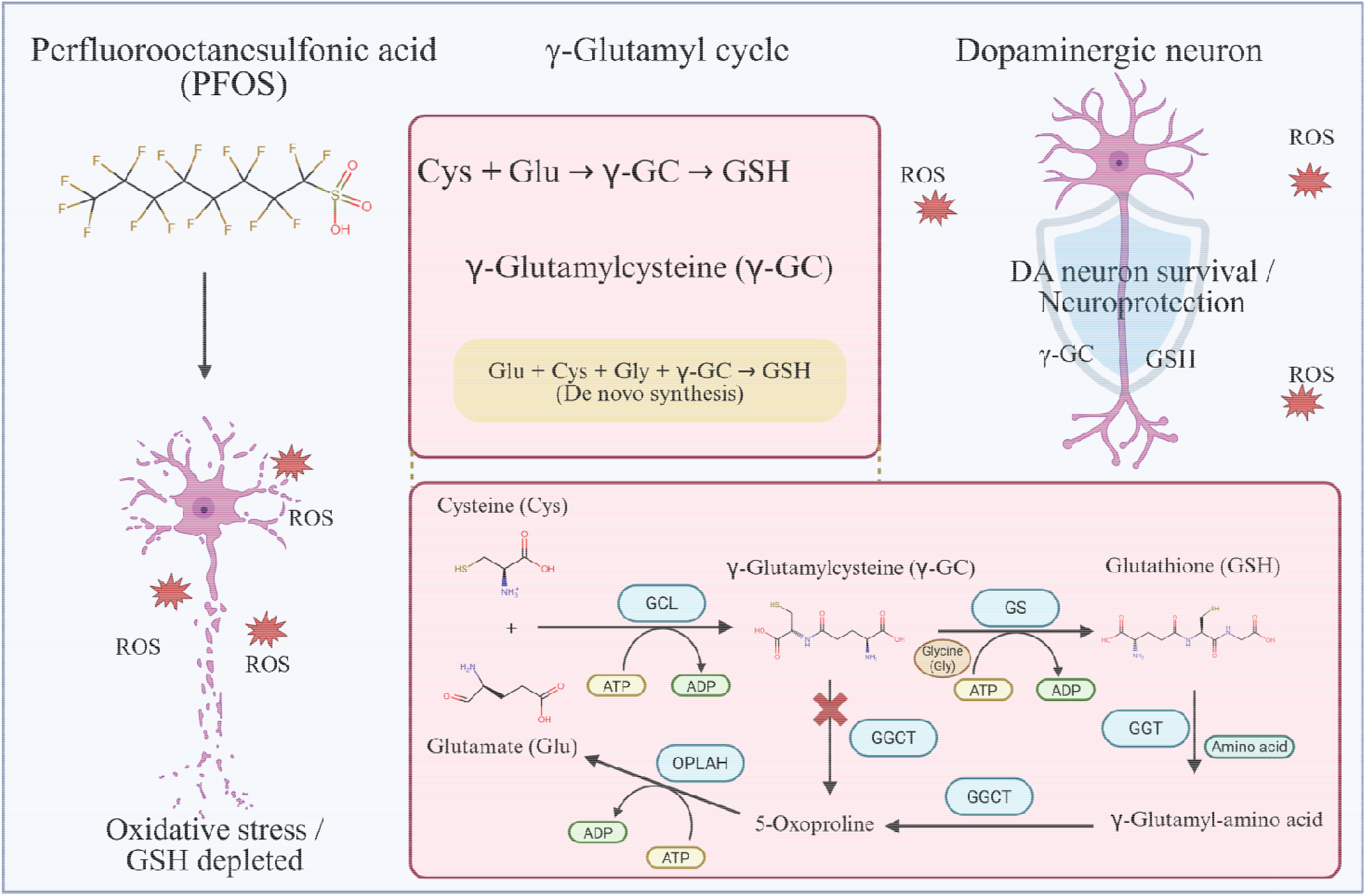
Mechanistic diagram: γ-Glutamyl-cysteine (γ-Glu-Cys) as a key intermediate in glutathione biosynthesis confers protection against PFOS-induced dopaminergic neurodegeneration. The glutathione pathway confers protection against PFOS-induced dopaminergic neurodegeneration. Exposure to perfluorooctane sulfonate (PFOS) results in dopaminergic (DA) neuron loss, as visualized in the BZ555 C. elegans strain expressing GFP in DA neurons. Cysteine and glutamate act as precursors for glutathione (GSH) synthesis. Co-treatment with GSH mitigates PFOS-induced DA neurodegeneration. GSH biosynthesis involves two enzymatic steps: first, glutamate and cysteine are ligated by glutamate-cysteine ligase (GCL) to form γ-glutamyl-cysteine (γ-Glu-Cys); then, glutathione synthetase (GS) adds glycine to produce GSH. Among the tested precursors, glutamate, cysteine, their combination, and γ-Glu-Cys, only γ-Glu-Cys significantly rescued DA neuron loss. As a key biosynthetic intermediate, γ-Glu-Cys plays a critical role in promoting GSH-mediated neuroprotection against PFOS toxicity. Excess γ-Glu-Cys is converted to γ-glutamyl-amino acid by the enzyme γ-glutamyl transferase (GGT). γ-Glutamyl-amino acid is converted to 5-oxoproline by the enzyme γ-glutamyl-cyclotransferase. 5-Oxoproline is next converted to glutamate by the enzyme 5-oxoprolinase (OPLAH). Glutamate is fed into the de novo synthesis pathway to generate GSH. Our studies identified that while γ-Glu-Cys is a key intermediate, 5-oxoproline does not confer neuroprotection. Rather, excess 5-oxoproline contributes to neurodegeneration, likely due to glutamate toxicity.

While the effect on Complex IV was less intense, it was observed at relatively lower concentrations of PFOS and eventually plateaued. We observed a significant inhibition of enzyme activity at doses, 5 ppm (0.864 ± 0.009, *p* < 0.0001), 12.5 ppm (0.789 ± 0.032, *p* < 0.0001), 25 ppm (0.718 ± 0.008, *p* < 0.0001), 50 ppm (0.709 ± 0.039, *p* < 0.0001), 125 ppm (0.715 ± 0.004, *p* < 0.0001) and 250 ppm (0.722 ± 0.032, *p* < 0.0001) as compared to control (1.000 ± 0.000) (Figure 6D). Overall, these studies identified that PFOS inhibits mitochondrial enzyme activity.

### GSH and γ-Glu-Cys rescue Complex III inhibition

In continuation to the observed beneficial effects of GSH and γ-Glu-Cys on dopaminergic cell loss, we questioned whether they can also rescue the complex III and complex IV enzyme inhibition. While neither GSH nor γ-Glu-Cys alleviated the complex IV enzyme inhibition, a significant rescuing effect of both GSH and γ-Glu-Cys was observed in complex III activity. In comparison to mitochondria exposed to PFOS 50 ppm (0.322 ± 0.082) an increase in normalized enzyme activity was observed upon co-exposure to 0.25 mM GSH (0.596 ± 0.037, *p* < 0.0001) and 0.25 mM γ-Glu-Cys (0.672 ± 0.034, *p* < 0.0001) (Figure 6E). Similarly, in mitochondria exposed to PFOS 75 ppm (0.214 ± 0.073), an increase in normalized enzyme activity was observed in mitochondria co-exposed to 0.25 mM GSH (0.439 ± 0.051, *p* = 0.0003) and 0.25 mM γ-Glu-Cys (0.467 ± 0.057, *p* = 0.0001) (Figure 6F).

## Discussion

Exposure to PFAS is a major public health and environmental concern[24,25]. PFOS is one of the most consistently detected PFAS in environmental and biological samples[5,26]. Our previous studies have discovered specific dopaminergic neurotoxicity and pathogenic pathways [7]. Preliminary results identified specific antioxidant pathways important to neuroprotection. NAC is an important precursor for GSH synthesis [27-30]. Given the lack of neuroprotective effect of NAC, we aimed to discover whether GSH itself is the sole neuroprotective agent in this pathway, or whether there may be a critical intermediate. Here, we have shown that γ-Glu-Cys is the critical intermediate. This study, for the first time, highlighted γ-Glu-Cys as a key intermediate in GSH-mediated alleviation of PFOS neurotoxicity, identifying a novel mechanism that fills the informational gap concerning the mechanism underlying PFOS neurotoxicity. The identification of specific neuroprotective pathways is critical to understanding PFAS neurotoxicity and the further development of treatment strategies.

Specifically, we tested whether the precursors of GSH biosynthesis, Cysteine, Glutamate, or combined, could alleviate neurotoxicity. Surprisingly, while cysteine provided slight neuroprotection, glutamate or cysteine plus glutamate were completely devoid of any neuroprotective effect. These findings, along with a lack of neuroprotective effect of NAC suggest that neuroprotection is not simply thiol mediated. This led us to further investigate the role of γ-Glu-Cys, the primary and highly conserved intermediate in GSH synthesis. Upon treatment, we observed significant neuroprotection within the dose range 0.125 to 0.5 mM. These results conclude that γ-Glu-Cys is a key intermediate in GSH-led neuroprotection. Additionally, in SH-SY5Y cells, we also observed a slight decrease in Total GSH levels; taken together, this suggests that GSH is a plausible target of PFOS toxicity. In our previous studies[7]. To investigate the effect on GSH synthesis pathway, we tested the effect on enzymes responsible for GSH synthesis, *gcs-1*/GCL, and *gss-1*/GS in *C. elegans* and SH-SY5Y cells. While only a slight decrease in mRNA expression of *gcs-1/*GCL was observed in *C. elegans*, no effect was observed on *gss-1*/GS. Paradoxically, a subtle, yet insignificant increase in the levels of GCL and GS was observed.

We further questioned if the observed neuroprotective effect could be due to alternative pathways. For example, unspent γ-Glu-Cys can be catalyzed by the enzyme, γ-GlutamylCycloTransferase to produce cysteine and 5-Oxoproline[20]. 5-Oxoproline (5-OP) is further catalyzed by the 5-oxoprolinase to synthesize glutamate[21]. Glutamate can be utilized to generate more GSH in combination with cysteine. However, glutamate is also known to exert neurotoxic effects, particularly through excitotoxicity [22,23]. Thus, it was important to first test whether 5-OP is involved in observed neurotoxicity. Upon testing higher doses, complete destruction of DA neurons was observed. Co-exposure studies using 0.125 to 0.5 mM 5-OP revealed cumulative neurotoxicity, ruling out the possibility of this alternative pathway conferring any neuroprotection. Furthermore, given that excess γ-Glu-Cys might result in the yield of 5-OP[20], suggests curtailed neuroprotection of γ-Glu-Cys at doses above 0.5 mM which was not the case with GSH that exhibits neuroprotection even at doses above 0.5 mM [7].

It is worth noting here that the previous studies identified mitochondria being primary target of PFOS neurotoxicity, exhibiting depletion in mitochondrial viability at doses far lower than neurotoxicity[7]. Given that, GSH is the principal antioxidant in mitochondria[10]. Even a subtle decrease in total GSH levels is likely to exacerbate mitochondrial GSH depletion since they are solely dependent on cytosolically synthesized GSH, which is not immediately available and relies on uptake through mitochondrial carrier proteins. Again, this study employing γ-Glu-Cys and 5-OP demonstrates an intricate, yet a delicate balance of chemicals with the ability to alter neurotoxicity both positively and negatively.

Based on cues from previous studies identifying mitochondria as primary targets [9], we further tested the effect of acute PFOS exposure on mitochondrial enzyme activity (Complexes I to IV). While no effect was observed on complex I, a significant inhibitory effect was observed on complexes II, III, and IV, supporting our hypothesis. Notably, complex III was more vulnerable in terms of extent of inhibition, whereas complex IV exhibited inhibition at far lower doses, though the extent of inhibition was modest. Overall, these results indicate that mitochondrial enzymes exhibit enzyme-specific and varied inhibition in response to PFOS exposure. Further, we tested whether GGC or GSH could rescue the enzyme inhibition. Using the same paradigm, we co-exposed the mitochondria to PFOS and 0.25 mM GSH and γ-Glu-Cys. It was found that both γ-Glu-Cys and GSH can alleviate complex III inhibition but cannot rescue complex IV inhibition, suggesting that the amelioration by γ-Glu-Cys and GSH cannot exert a protective effect downstream of complex III. γ-Glu-Cys alone has also been reported to confer protection independent of GSH synthesis through disposal of peroxide radicals and acting as a cofactor for glutathione peroxidase-1[31], which is known to express in mitochondria[32,33]. One caveat of our study was that the direct effect of GSH on the enzyme GCL could not be studied due to the purified enzyme/reliable assays for GCL being unavailable. Future studies will investigate the mechanisms related to mitochondrial GSH uptake. Overall, this study has identified a critical intermediate in GSH-mediated neuroprotection against PFOS-induced neurotoxicity.

## Author contributions

SRS and JRC helped with funding and conceptualization of studies. In addition, SRS performed studies related to C. elegans, SH-SY5Y, and mitochondrial enzyme activity. NB performed studies related to GSH estimation and mitochondrial enzyme activity. NCK helped with technical support related to C. elegans culture. This work was supported by the National Institute of Environmental Health Sciences.

## Declarations

### Ethical approval and consent to participate

All animal experiments conducted were in compliance to Institutional IACUC.

### Data availability

The data that supports the findings of this study are available from the author upon reasonable request.

## Acknowledgement

Strains were provided by the CGC, which is funded by NIH Office of Research Infrastructure Programs (P40 OD010440).

